# Effects of early life adversity on meningeal mast cells and proinflammatory gene expression in male and female *Mus musculus*

**DOI:** 10.1101/2021.09.17.460793

**Authors:** Natalia Duque-Wilckens, Erika Sarno, Robby E. Teis, Frauke Stoelting, Sonia Khalid, Zakaria Dairi, Alex Douma, Nidia Maradiaga, Kyan Thelen, A.J. Robison, Adam J. Moeser

**Affiliations:** Department of Physiology, Michigan State University, East Lansing, MI, 48824; Department of Large Animal Clinical Sciences, College of Veterinary Medicine, Michigan State University, East Lansing, MI, 48824

**Author notes:** To whom correspondence should be addressed: Gastrointestinal Stress Biology Laboratory, Department of Large Animal Clinical Sciences, College of Veterinary Medicine, Michigan State University, East Lansing, MI 48824.

## Abstract

Exposure to early life adversity (ELA) in the form of physical and/or psychological abuse or neglect increases the risk of developing psychiatric and inflammatory disorders later in life. It has been hypothesized that exposure to ELA results in persistent, low grade inflammation that leads to increased disease susceptibility by amplifying the crosstalk between stress-processing brain networks and the immune system, but the mechanisms remain largely unexplored. The meninges, a layer of three overlapping membranes that surround the central nervous system (CNS)- duramater, arachnoid, and piamater – possess unique features that allow them to play a key role in coordinating immune trafficking between the brain and the peripheral immune system. These include a network of lymphatic vessels that carry cerebrospinal fluid from the brain to the deep cervical lymph nodes, fenestrated blood vessels that allow the passage of molecules from blood to the CNS, and a rich population of resident mast cells, master regulators of the immune system. Using a mouse model of ELA consisting of neonatal maternal separation plus early weaning (NMSEW), we sought to explore the effects of ELA on duramater mast cell histology and expression of inflammatory markers in male and female C57Bl/6 mice. We found that mast cell number, activation level, and relative expression of pseudopodia differ across duramater regions, and that NMSEW exerts region-specific effects on mast cells in males and females. Using gene expression analyses, we next found that NMSEW increases the expression of inflammatory markers in the duramater of females but not males, and that this is prevented by pharmacological inhibition of mast cells with ketotifen. Together, our results show that ELA drives sex-specific, long-lasting effects on the duramater mast cell population and immune-related gene expression, suggesting that the long-lasting effects of ELA on disease susceptibility could be partly mediated by meningeal function.

## INTRODUCTION

Exposure to early life adversity (ELA), in the form of physical and/or psychological abuse or neglect, increases the risk of developing disorders at a multisystemic level later in life^1–4^: children with history of ELA show higher rates of psychiatric diseases such as unipolar depression and substance abuse^5,6^, but also increased prevalence of gastrointestinal disorders^7,8^, multiple sclerosis^9^, migraines^10^, cardiovascular disease^11^, and metabolic syndrome^12^,among others. Since these disorders share a mechanistic origin involving inflammation^13–18^, it has been proposed that ELA-driven changes in the immune system are the root underlying susceptibility to these diseases. Indeed, children^19,20^ and adults^21– 23^ with history of ELA display elevated levels of inflammatory markers. This, together with findings that ELA increases sensitivity of brain circuits involved in threat processing^24–26^, prompted the hypothesis ELA drives persistent systemic inflammatory states by amplifying the crosstalk between stress-processing brain networks and the immune system^27,28^.

A crucial interface between the brain and the peripheral immune system is the meninges, a layer of three overlapping membranes - duramater, arachnoid, and piamater - that surround the central nervous system (CNS). Although originally thought to serve mostly as a physical protection, there is growing evidence the meninges play an active role in coordinating immune trafficking throughout the CNS^29^ through a variety of mechanisms. First, vessels within the duramater, unlike cerebral vessels, are fenestrated, and thus open to the passage of molecules present in the blood^30^. Second, the meninges hold vascular channels connecting the skull bone marrow with the brain surface, allowing myeloid cell migration into the brain^31^. Third, the duramater provides a network of lymphatic vessels that carry cerebrospinal fluid from the brain to the deep cervical lymph nodes^32–35^, which permits cerebrospinal-injected immune cells to traffic from the brain to the periphery. Finally, the meninges contain a rich population of resident immune cells, among which are mast cells^36^, master regulators of both innate and adaptive immune responses^37^.

Mast cells are multifunctional innate immune cells characterized by their biphasic response to stimuli, which involves an early release of a wide range of preformed granule mediators (e.g. histamine, serotonin, proteases) followed by release of *de novo* synthesized mediators (e.g. cytokines). These mediators allow them to not only orchestrate innate and adaptive immune responses^37^, but also modulate activity of vascular and lymphatic endothelial cells^38^, nerves^39^, and glia^40,41^. Meningeal mast cells, specifically, are key in maintaining blood brain barrier integrity^42,43^ and recruiting T-cells and neutrophils^44^. Menigeal mast cells can also mediate the inflammatory responses associated with brain stroke^42,45^ and the activation of the trigeminal pain pathway associated with migraines^46^. Importantly, mast cells can rapidly respond to stress^47–49^, and ELA can persistently modify their activity and distribution^50^. Together, this suggests that meningeal mast cells are ideally positioned to play a key role in the brain-immune crosstalk driving persistent inflammation associated with ELA, but, to our knowledge, whether ELA has long-lasting effects on meningeal mast cell biology remains undetermined.

Here, we assessed the effects of ELA on duramater mast cell histology and expression of inflammatory markers with and without exposure to adult stress in male and female C57Bl/6 mice. Our analyses included mouse interleukin 6 (IL-6) and mouse tumor necrosis factor alpha (TNF-alfa), major proinflammatory cytokines associated with ELA-related diseases^51,52^; mouse vascular endothelial growth factor C (VEGF-C), critical for meningeal lymphangiogenesis^53^ and recently shown to be actively involved in brain immunosurveillance^54^; as well as mouse chymotryptic serine proteinase (CMA1), mouse histidine decarboxylase (Hdc), and mouse tryptophan hydroxylase 1 (Tph1), which reflect different pathways of mast cell activity^55–57^. We used the model of neonatal maternal separation plus early weaning (NMSEW). Previous work has demonstrated that adult animals exposed to neonatal maternal separation show elevated peripheral^58–61^ and central^62–64^ inflammatory markers, are more susceptible to develop gastrointestinal^65,66^, autoimmune^67^ and cardiorespiratory^68,69^ abnormalities, and show increased anxiety and depressive-like behaviors^58,70,71^, making this an ideal model to assess whether meningeal immunity plays a role in ELA-induced persistent systemic inflammation. We added an early weaning component to this protocol to increase the likelihood of inducing robust, long-lasting phenotypes^72^. Finally, to further assess whether mast cell activity plays a role in ELA-induced changes in meningeal biology, we administered the mast cell inhibitor Ketotifen fumarate (Ketotifen) before exposure to adult stress. We hypothesized that NMSEW would increase activity of mast cells and expression of inflammatory markers in the duramater after exposure to adult stress, and that this would be prevented by administration of Ketotifen.

## MATERIALS AND METHODS

### Animals

All animal procedures were performed in accordance with the regulations of the Michigan State University animal care committee. Founding C57Bl/6 breeders were obtained from the Jackson Laboratory (Bar Harbor, Me). All animals were kept in cages enriched with Nestlet and Bed-r’Nest**^®^** at 20–23°C under a 12 h light/dark cycle with ad libitum access to food and water and housed in same sex groups of 3-5 individuals or in adult male-female pairs for breeding purposes (male was removed from the cage before litters were born). For all experiments, newborn litters were randomly assigned to one of two conditions: neonatal maternal separation + early weaning stress (NMSEW) or normally handled (NH). For experiment 1, all animals from all litters were exposed to adult stress. For all gene expression analyses (experiments 2,3 and 4), each litter was divided and randomly assigned to one of the three experiments to minimize litter effects.

### Neonatal maternal separation plus early weaning (NMSEW)

During postnatal (PN) days 1-17, pups in the NMSEW group were daily separated from their dams and placed in individual containers for 180 minutes, after which they were returned to their home cage with their dam and siblings. They were left undisturbed between separation events. On day 17, NMSEW pups were weaned and housed in same sex sibling groups. HydroGel™ was provided for the first two days after weaning to ensure proper hydration. Pups in the NH groups were left undisturbed until postnatal day 28, when they were weaned. All animals were ear punched for individual identification on weaning day. After weaning, mice from NMSEW and NH groups were left undisturbed until they reached 8 weeks old (PN 56, adult) **(Fig. 2A,3A)**.

**Fig. 1.**
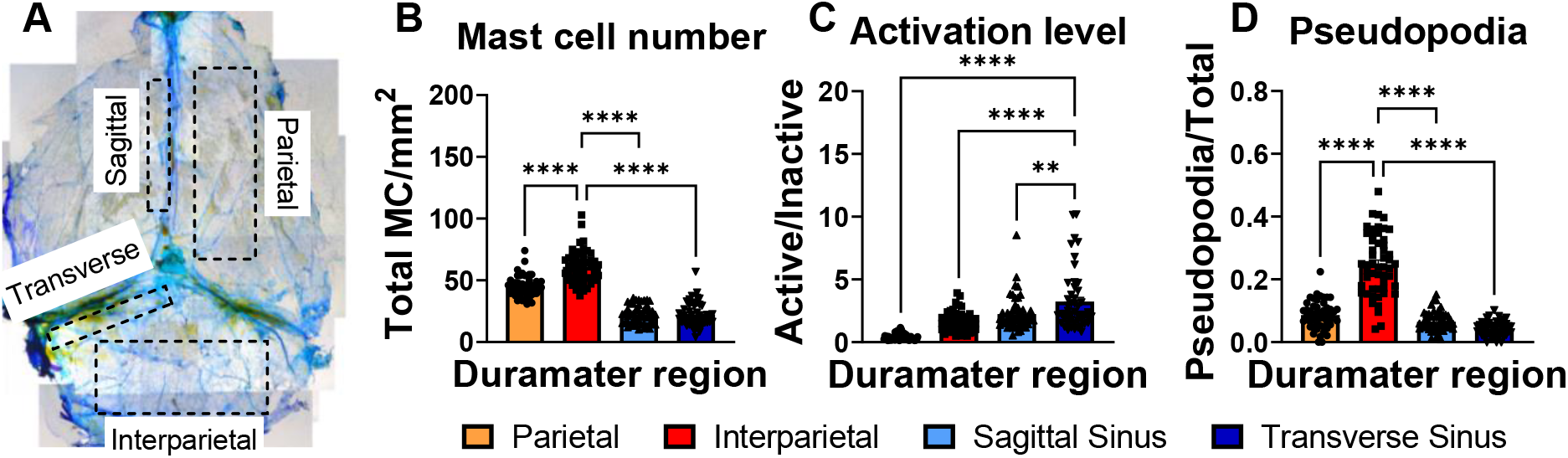
Mast cells are not homogeneous across duramater regions. A: diagram showing regions of duramater analyzed:: parietal, interparietal, Sagittal sinus and Transverse sinus. B: Total mast cell (MC) number per mm^2^ significantly differs by duramater region, with interparietal region showing the highest mast cell density. C: Activation level of mast cells differs by duramater region, with transverse region showing the highest activation. D: Relative expression of pseudopodia varies by duramater region, with interparietal region showing the highest expression. One way ANOVA followed by Dunnett’s multiple comparisons test, **p<0.01, ****p<0.0001. n=49-52 per group.

**Fig. 2.**
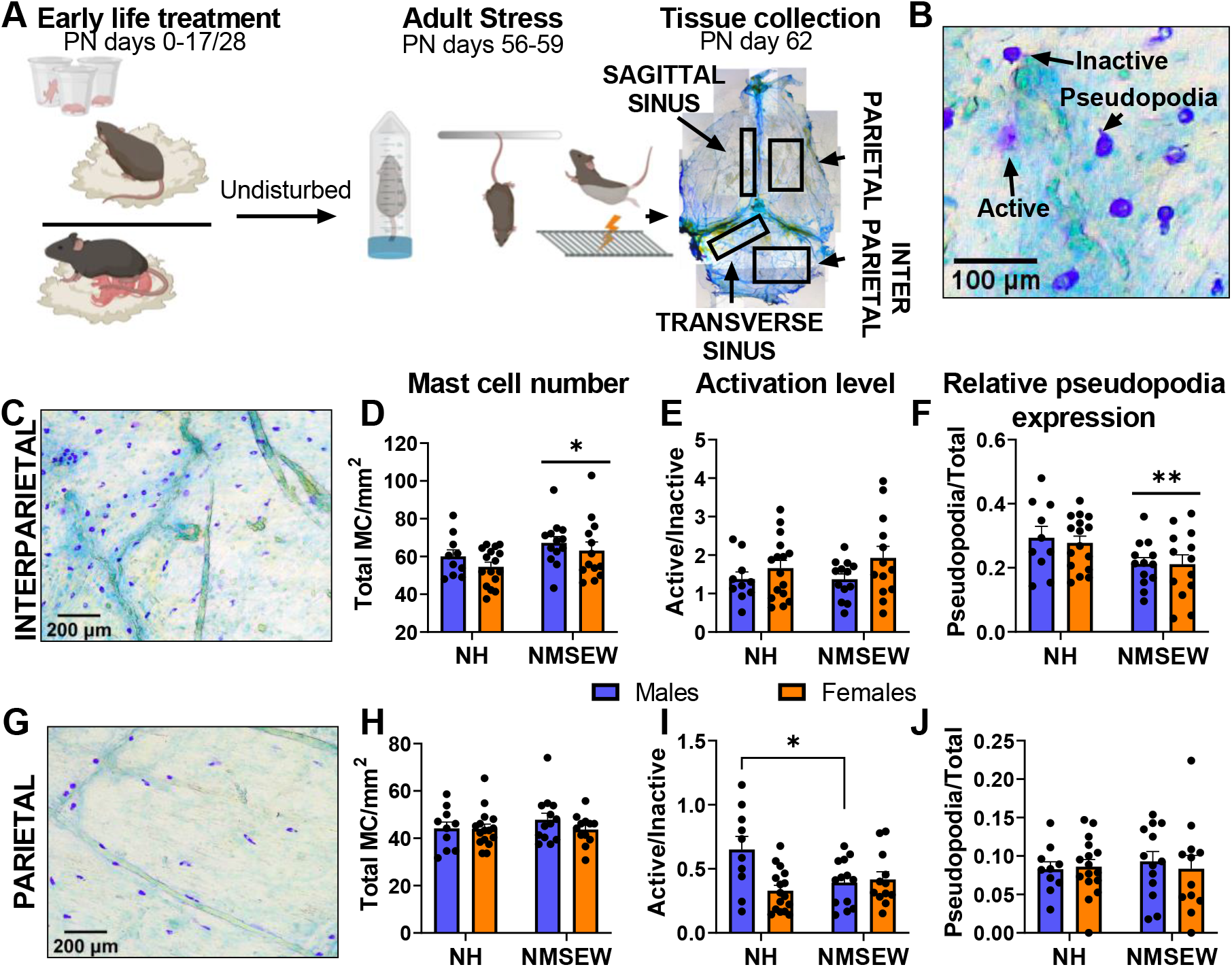
Early life adversity exerts long-term effects on duramater mast cells in interparietal and parietal regions of the duramater. **A:** diagram showing experimental timeline. **B**. Photomicrograph showing examples of mast cell features quantified: active mast cell, inactive mast cell, and presence of pseudopodia. **C, G**. Photomicrograph showing examples of mast cells in interparietal and parietal regions, respectively. **D**. Compared to NH, NMSEW animals show higher numbers of mast cells in interparietal region (two way ANOVA main effect of early life treatment, *p=0.03). **F**. Compared to NH, NMSEW animals show lower relative expression of pseudopodia (two way ANOVA main effect of early life treatment **p=0.006). **I**. In the parietal region, NMSEW reduced the activation levels in males (Šídák’s multiple comparisons test*p=0.01). No differences between groups were seen in **E**. interparietal activation levels of mast cells, **H**. parietal total mast cell number, and **J**. Parietal relative expression of pseudopodia. N= 10-16 per group.

**Fig. 3.**
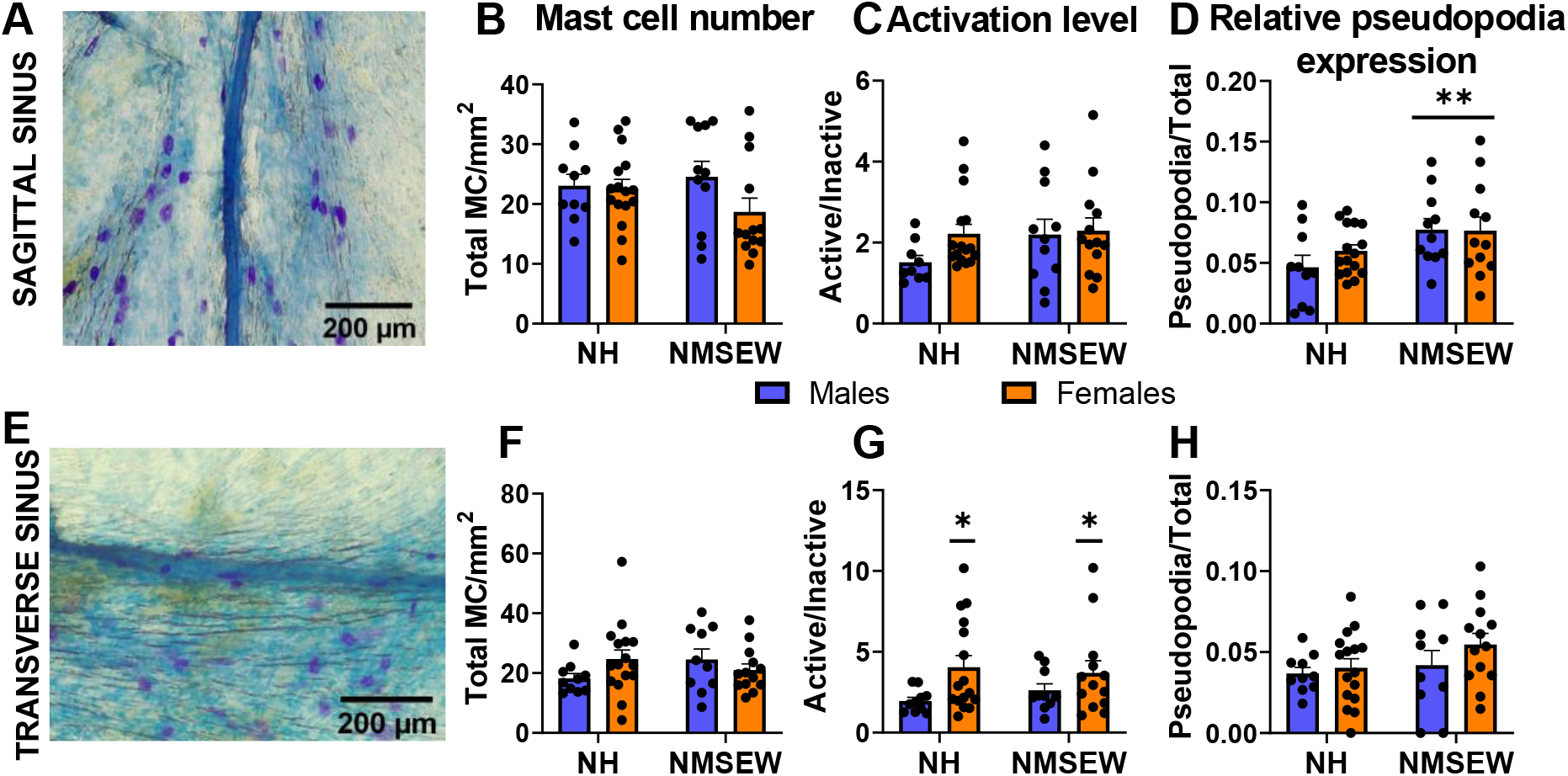
Early life adversity exerts long-term effects on duramater mast cells in sagittal and transverse sinus regions of the duramater. **A, B**. Photomicrograph showing examples of mast cells in sagittal and transverse regions, respectively. **D**. Compared to NH, NMSEW animals show higher numbers of relative pseudopodia in the sagittal sinus region (two way ANOVA main effect of early life treatment, **p=0.008). **G**. In the transverse sinus region, females showed higher activation levels than males (two way ANOVA main effect of sex, *p=0.02). No differences between groups were seen in **B,F**. Mast cell number in sagittal or transverse sinus regions, **C**. Mast cell activation level in sagittal sinus region, or **H**. relative expression of pseudopodia in transverse sinus region. N= 10-16 per group.

### Adult Stress

The adult stress consisted of a shortened version of subchronic variable stress^73,74^: for three consecutive days, adult animals were singly housed and exposed for 10 min to either tail suspension test, restraint stress, or foot shock stress (20 total 0.45 mA shocks at random). This milder version of stress was chosen to minimize ceiling effects based on evidence showing that early life adversity increases sensitivity to future stress^75^. **In experiments 1 and 3** (Figs.2A,3A and 5A), all animals from both the NMSEW and NH groups were exposed to adult stress on days 56-59. **In experiment 2** (Fig.4A), animals from the NMSEW and NH groups were left undisturbed until tissue collection. **In experiment 4** (fig.6A), animals from both the NMSEW and NH groups received an intraperitoneal injection of Ketotifen fumarate (see below) before each episode of adult stress on days 56-59.

**Fig. 4.**
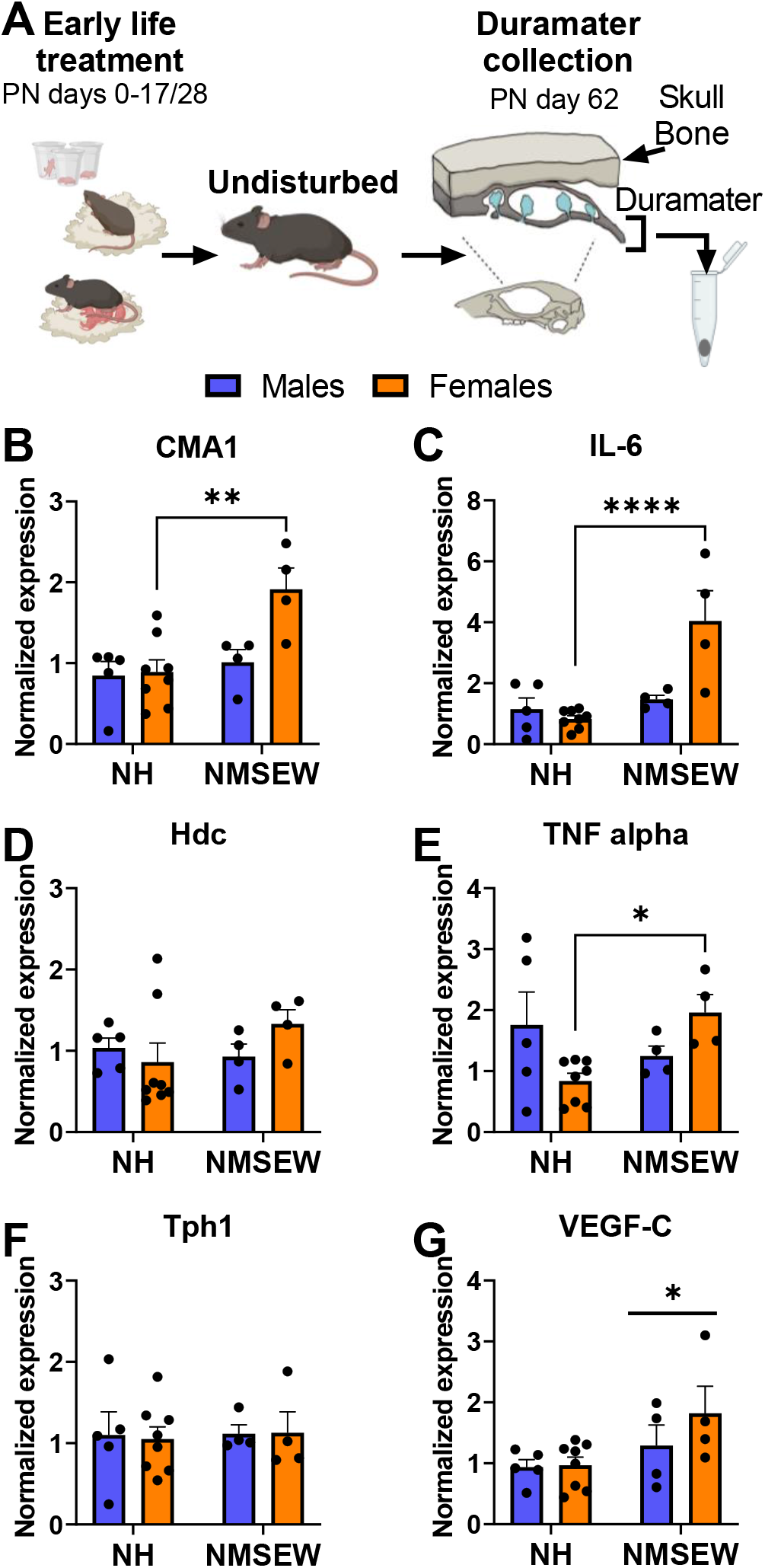
NMSEW increases duramater expression of proinflammatory molecules in females but not males. **A**. Experimental timeline. **B, C, E**. NMSEW increased expression of CMA1 (Šídák’s multiple comparisons test **p=0.002) IL-6 (Šídák’s multiple comparisons test ****p<0.0001), and TNF alpha (Šídák’s multiple comparisons test *p=0.03) in females but not males **G**. The expression of VEGF-C was increased in animals exposed to NMSEW compared to NH (two way ANOVA main effect of early life treatment *p=0.026). No effects of sex or early life treatment where seen in **D**. Hdc, or **F**. Tph1. N= 4-8 per group.

**Fig. 5.**
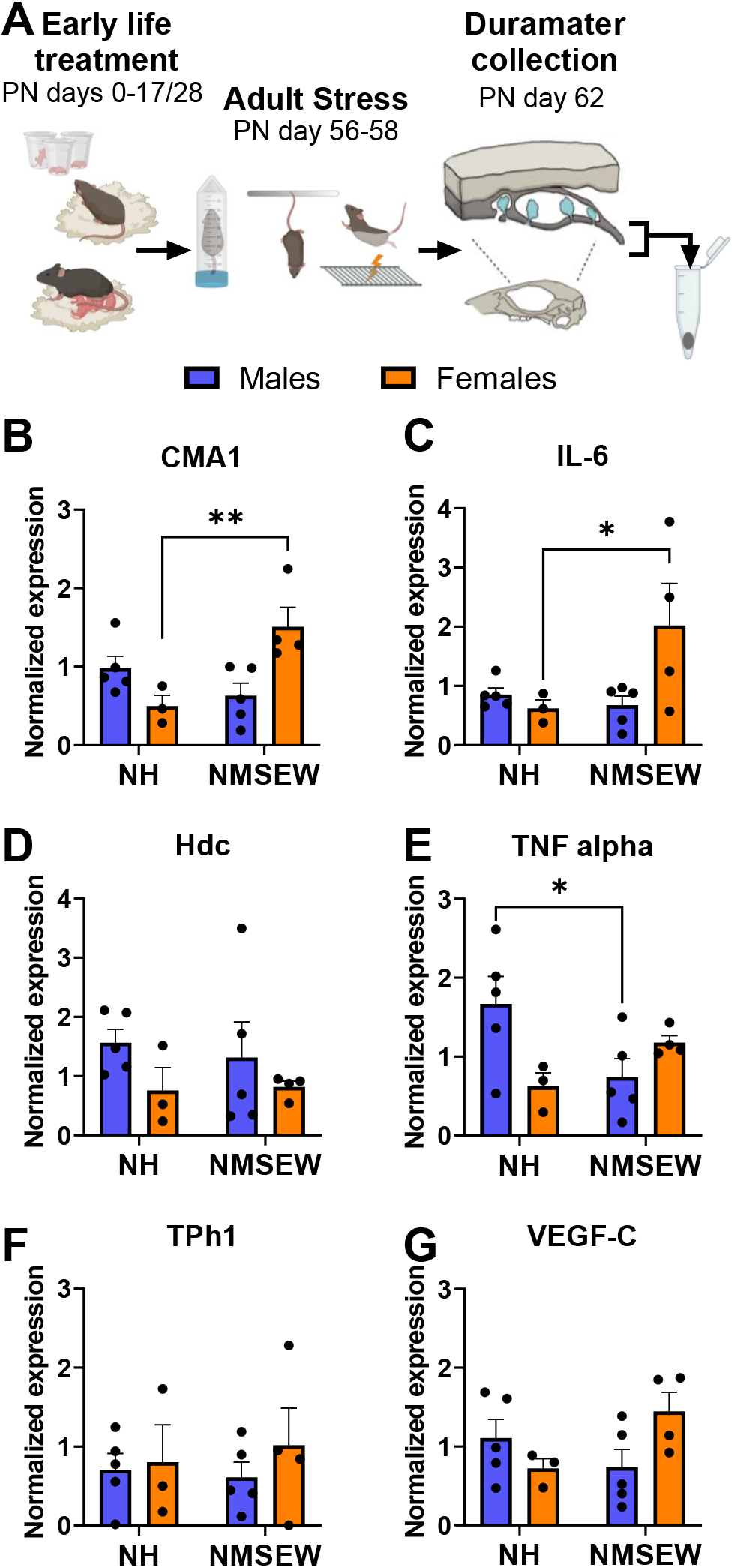
Exposure to a second hit stress doesn’t interfere with the effects of NMSEW on meningeal gene expression. **A**. Experimental timeline. **B, C**. NMSEW increased expression of CMA1 (Šídák’s multiple comparisons test **p=0.007) and IL-6 (Šídák’s multiple comparisons test *p=0.049) in females but not males. **E**. NMSEW reduced the expression of TNF alpha in males (Šídák’s multiple comparisons test *p=0.034). No effects of sex or early life treatment where seen in **D**. Hdc, **F**. Tph1, of **G**. VEGF-C. N= 3-5 per group

**Fig. 6.**
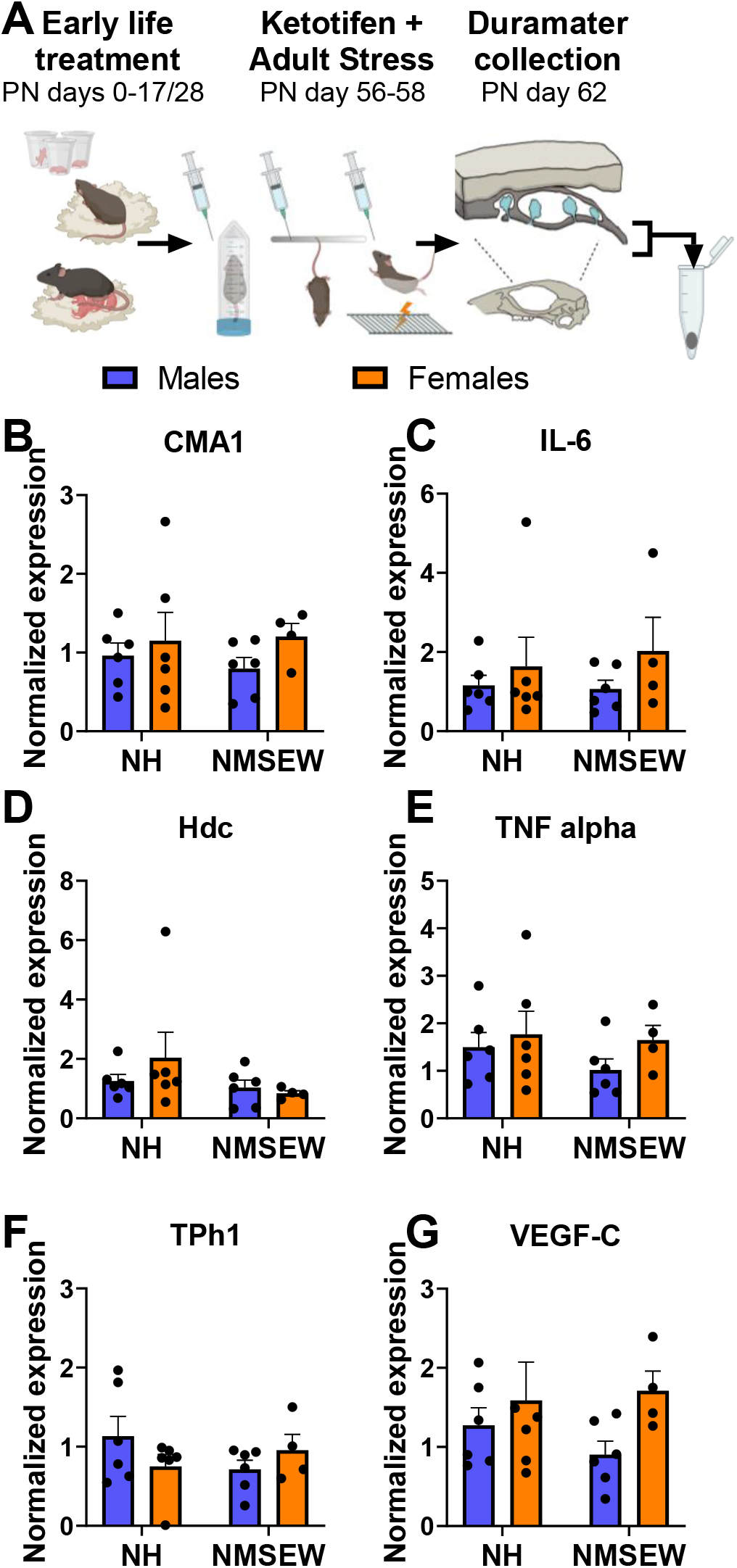
Stabilizing mast cells before exposure to adult stress eliminates the effects of NMSEW on meningeal gene expression. **A**. Experimental timeline. B,C,D,E,F,G. No effects of early life treatment or sex were seen in the expression of any of the genes measured after administration of Ketotifen.

### Mast cell stabilization

20mg/kg Ketotifen (Cayman Chemical, Catalog #20303) was administered intraperitoneally 30 min prior each episode of adult stress (fig.5A). Ketotifen fumarate is a histamine H1 receptor antagonist and mast cell stabilizer (prevents mast cell degranulation)^76,77^ that crosses the blood brain barrier^78^. The chemical structure of ketotifen fumarate is available in PubChem, PubChem CID: 5282408. 20 mg/kg dose was based on previously published studies showing successful mast cell stabilization in mice^79–81^.

### Tissue collection

3 days after the last episode of stress, animals were euthanized via cervical dislocation. **In experiment 1**, the skull caps with attached duramater were collected and immediately placed into 10% formalin solution. **In experiments 2, 3, and 4**, the duramater was freshly separated from the skull and immediately flash frozen using dry ice.

### Duramater mast cell staining and quantification (experiment 1)

After 24h of postfixation with 10% formalin, duramater was carefully separated from the skull, washed with DI water, and mounted on Superfrost plus microscope slides (Fisher Scientific). Once dry, slides were immersed for 1h in a solution containing 0.5% Toloudine blue in 0.5N Hydrochloric acid, after which they were washed in DI water, dried, and coverslipped using Vectashield mounting medium. A Leica DM750 bright field Microscope with ICC50W Integrated Camera was used to take four 10x brightfield pictures per each duramater region available (fig.2A,3A): parietal, interparietal, sagittal sinus, and transverse sinus. Finally, quantification was made by a blind experimenter using ImageJ. A 1000×780 μm box was used to limit counting areas:two random and non-overlapping counting areas were used per picture, 8 counting areas total per duramater region. The boxes for parietal and interparietal regions were placed parallel to the sinuses, so that the sinus blood vessel and lumen were not included in the counting region. The total number of mast cells, number of active mast cells, number of inactive mast cells, and number of mast cells expressing pseudopodia were quantified per counting area. For detailed description of criteria used to classify mast cells please refer to table 1. The level of activation was calculated as the number of active mast cells/number of inactive mast cells, and the relative expression of pseudopodia was calculated as number of mast cells expressing pseudopodia/total number of mast cells per counting area. The data are presented as average numbers per area/mouse (total counts/number of counting areas) **(fig.2B,3B)**.

**Table 1.**
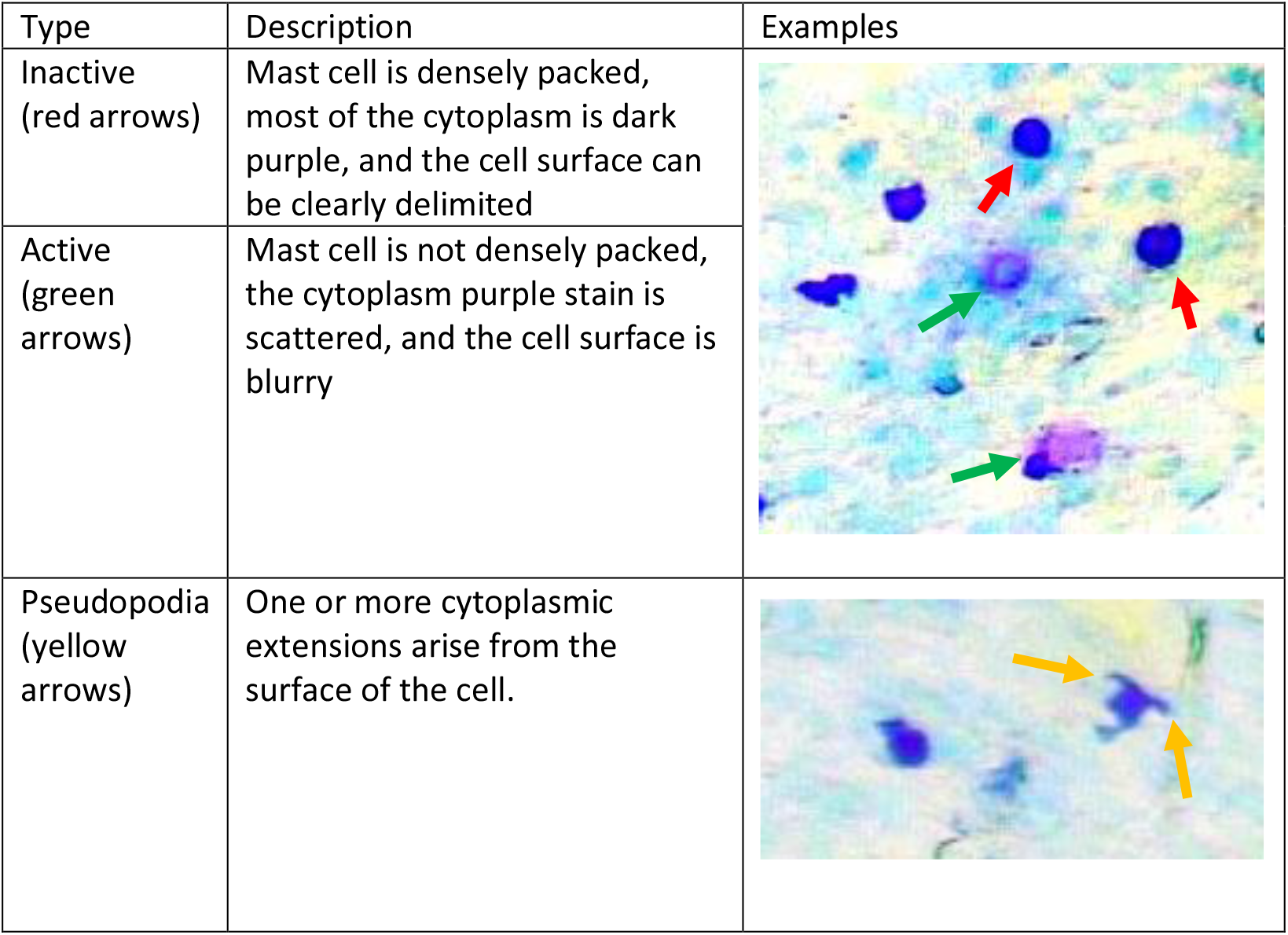
Classification of duramater mast cells.

| Type | Description | Examples |
| --- | --- | --- |
| Inactive (red arrows) | Mast cell is densely packed, most of the cytoplasm is dark purple, and the cell surface can be clearly delimited |  |
| Active (green arrows) | Mast cell is not densely packed, the cytoplasm purple stain is scattered, and the cell surface is blurry |  |
| Pseudopodia (yellow arrows) | One or more cytoplasmic extensions arise from the surface of the cell. |  |

**Table 1. Classification of duramater mast cells**
| Type | Description | Examples |
| --- | --- | --- |
| Inactive<br>(red arrows) | Mast cell is densely packed, most of the cytoplasm is dark purple, and the cell surface can be clearly delimited |  |
| Active<br>(green arrows) | Mast cell is not densely packed, the cytoplasm purple stain is scattered, and the cell surface is blurry |  |
| Pseudopodia<br>(yellow arrows) | One or more cytoplasmic extensions arise from the surface of the cell. |  |

### Gene expression analyses (experiments 2, 3, and 4)

Duramater was homogenized using Trizol (Invitrogen), and total RNA was extracted using the RNAeasy MicroKit-Qiagen #74004. RNA integrity and concentration were assessed using an Agilent 2100 bioanalyzer (Agilent Technologies, Palo Alto, California). Samples with RIN values equal to or above 8 were converted to cDNA using the Applied Biosystems™ High-Capacity cDNA Reverse Transcription Kit (#4368814). The cDNA was used on the real-time RT^2^ Profiler PCR Array (QIAGEN, Cat. no. CLAM38697) in combination with RT^2^ SYBR^®^Green qPCR Mastermix (Cat. no. 330503). Each array plate contained one set of 96 wells with a panel of Qiagen designed primers for mouse IL-6, TNF-alfa, Hdc, Tph1, VEGF-C, and CMA1. Positive PCR controls were included in each 96-well set on each plate. Glyceraldehyde-3-phosphate-dehydrogenase (*GAPDH*) was used as the assay reference gene. Each array contained samples from control and test groups.

### Statistical analyses

All figures and statistical analyses were done using Graphpad Prism 9 software. For all analyses, we used an alpha level criterion of 0.05 for statistical significance. One-way ANOVAs followed by Dunnett’s multiple comparison tests were used to compare mast cells between duramater regions (fig 1). Two-way ANOVAs were used to analyze effects of sex, early life treatment, and the interaction in all experiments (figs.2-6), followed by Šídák’s multiple comparisons test in ANOVAS showing significant interaction effects.

## RESULTS

### Mast cells are not homogeneous across duramater regions

Since the cellular compartment of the duramater is not homogeneous^29^, we first assessed whether there were overall differences in mast cell number, activation levels, and pseudopodia expression among the major regions of the cortical duramater: interparietal, parietal, sagittal sinus, and transverse sinus. We found that the region of the duramater had a significant effect on all the measures. There was a main effect of region on the average of mast cells per mm^2^ (one way ANOVA F3,198=185.5, p<0.0001, Fig. 1B), with the interparietal duramater showing significantly higher mast cell density than any other region (Dunnett’s multiple comparisons test interparietal vs. parietal p<0.0001, interparietal vs sagittal p<0.0001, interparietal vs. transverse sinus p<0.0001). Similarly, there was a significant effect of duramater region on activation level (one way ANOVA F3,198=34.96, p<0.0001, Fig. 1C), with transverse sinus showing the highest ratio of active to inactive mast cells (Dunnett’s multiple comparisons test transverse vs. parietal p<0.0001, transverse vs interparietal p<0.0001, transverse vs. sagittal sinus p<0.01). Finally, the relative expression of pseudopodia, cytoplasmic extensions that mast cells use to transfer granule content to neighboring cells^82^ and/or engulf pathogenic cells^83^ was also different between duramater regions (one way ANOVA F3,198=138.8, p<0.0001, Fig. 1D), with the interparietal duramater showing the highest relative expression of pseudopodia (Dunnett’s multiple comparisons test interparietal vs. parietal p<0.0001, interparietal vs sagittal p<0.0001, interparietal vs. transverse sinus p<0.0001).

### NMSEW induces changes in duramater mast cells in a region-specific fashion

Because our results demonstrate heterogeneous characteristics of meningeal mast cells depending on the region of the duramater, all subsequent comparisons between treatments were done within specific duramater regions. In the **interparietal** region of the duramater, early life treatment affected total mast cell number and relative pseudopodia expression. Compared to NH animals, NMSEW animals showed significantly higher mast cell number (two way ANOVA main effect of early life treatment F(1,48)=5.1, p=0.03, **fig. 2D**) and lower relative expression of pseudopodia (two way ANOVA main effect of early life treatment F(1,48)=8.2, p=0.006, **fig. 2F**). No differences were seen between groups in the activation levels.

In the **parietal** duramater, there were no differences between groups in the mast cell number or pseudopodia expression, but there was an interaction between early life treatment and sex in the activation levels (two way ANOVA interaction F(1,47)=7.9, p=0.007): in males but not females, NMSEW reduced the activation level (Šídák’s multiple comparisons test p=0.01, **fig. 2I**). In the **sagittal sinus** region, the treatment during early life also affected the relative expression of pseudopodia, but, in contrast to the parietal region, NMSEW animals showed higher pseudopodia expression (two way ANOVA main effect of early life treatment F(1,48)=7.5, p=0.008, **fig. 3F**). There were no differences between groups in the total number or activation levels of mast cells.

Finally, in the **transverse sinus** region, while no effects of early life treatment were found in any of the measures, there was an effect of sex on activation levels, with females showing higher activation levels than males (two way ANOVA main effect of sex F(1,45)=5.74, p=0.02, **fig. 3I**).

### NMSEW increases duramater expression of proinflammatory molecules in females but not males

There was a significant interaction between early life treatment and sex in expression of CMA1 (two way ANOVA interaction effect F(1,17)=5.008, p=0.04, **fig. 4B**), IL-6 (two way ANOVA interaction effect F(1,17)=11.2, p=0.003, **fig. 4C**), and TNF alpha (two way ANOVA interaction effect F(1,17)=6.76, p=0.02, fig. **4E**). Šídák’s multiple comparisons test revealed that NMSEW increased the expression of all three genes in females (CMA1 p=0.002, IL-6 p<0.0001, TNF alpha p=0.03) but not in males. The expression of VEGF-C was increased in animals exposed to NMSEW compared to NH (two way ANOVA main effect of early life treatment F(1,17)=6, p=0.026, **fig. 4G**). We found no differences in the expression of Hdc or Tph1 between groups.

### Exposure to a second hit stress doesn’t interfere with the effects of NMSEW on meningeal gene expression

Similar to experiment 2, in experiment 3 there was a significant interaction between early life treatment and sex in expression of CMA1 (two way ANOVA interaction effect F(1,13)=13.34, p=0.003, **fig. 5B**), IL-6 (two way ANOVA interaction effect F(1,13)=1.48, p=0.04, **fig. 5C**), and TNF alpha (two way ANOVA interaction effect F(1,13)=7.7, p=0.02, **fig. 5E**). Šídák’s multiple comparisons test revealed that NMSEW increased the expression of CMA1 (p=0.007) and IL-6 (p=0.049) in females. NMSEW reduced the expression of TNF alpha in males (p=0.034), while the effect in females was not significant. There was also a significant interaction in the expression of VEGF-C (two way ANOVA interaction effect F(1,13)=5.4, p=0.036, **fig. 5G**), but no significant differences between groups were apparent after Šídák’s multiple comparisons test. No effects of early life treatment or sex were found in the expression of Hdc or Tph1.

### Stabilizing mast cells before exposure to adult stress eliminates the effects of NMSEW on meningeal gene expression

In animals receiving Ketotifen, no effects of early life treatment or sex were seen in the expression of any of the genes measured (fig. **6 B,C,D,E,F,G**). These data indicate that mast cell activity is critical for the effects of NMSEW on meningeal gene expression.

## DISCUSSION

In the present study we show that mast cell number, activation level, and relative expression of pseudopodia differ across duramater regions, and that NMSEW exerts meningeal region-specific effects on mast cells in males and females. Using gene expression analyses, we next showed that NMSEW increases the expression of inflammatory markers in the duramater of females but not males, and that this is prevented by pharmacological inhibition of mast cells with ketotifen. Together, our results show that early life adversity results in sex-specific, long-lasting effects on meningeal mast cell population and immune-related gene expression and suggest that that these are at least partly driven by changes in mast cell activity.

### Early life adversity exerts long-term effects on duramater mast cells in a region-specific fashion

Using T-blue staining in whole mount duramater from adult animals, we found that mast cell populations are not homogeneous across duramater regions. Compared to parietal, sagittal sinus, and transverse sinus, the interparietal region showed the highest number of mast cells per mm^2^ and the highest relative expression of pseudopodia. Conversely, the transverse sinus showed the highest activation levels. Based on these results and previous evidence showing that the duramater has location-specific mechanical and biochemical properties^84^, innervation^85,86^, presence of blood and lymphatic vessels^32,33^, as well as cellular immunity^87^, we separated the mast cell histology analyses by duramater region.

In the **interparietal region** of the duramater, NMSEW increased the total mast cell number while decreasing the relative activation levels, regardless of sex. This region of the duramater overlays the cerebellum, a brain region involved in motor coordination but also in cognitive and emotional processes ^88–90^. Interestingly, human and animal studies have shown that ELA has long-term structural and functional consequences on the cerebellum^91^. For example, physical/psychological neglect during early life can affect cerebellar volume in healthy children^92^ and adults suffering from obsessive compulsive disorder^93^. In rats, NMSEW is associated with reduced cerebellar metabolic activity^94^. Although it is unknown whether these effects of ELA on cerebellum are mediated by interparietal meninges, some evidence suggest that this could be possible: the interparietal meninges are directly involved in the migration of cells from the external germinal layer into the internal granule layer in the cerebellum during both embryonic and postpartum period^95,96^. Future studies assessing the role of interparietal mast cells in cerebellar function, and whether this is affected acutely and/or chronically by ELA, could reveal a novel communication pathway between the meninges and the brain.

In the **sagittal sinus region**, NMSEW increased the relative expression of mast cell pseudopodia regardless of sex. Pseudopodia represent a mechanism by which mast cells directly communicate with adjacent cells in a process called transgranulation, in which mast cells directly transfer their granular content to other cells^82,97^. The meningeal lymphatic network runs along the sagittal and transverse sinus regions ^32,33^. This network is critical for the cross-talk between the CNS and the peripheral immune system, as it provides an exclusive route for draining macromolecules and trafficking immune cells from the cerebrospinal fluid to the cervical lymph nodes^32,98,99^. Interestingly, NMSEW also increased the expression of VEGF-C, the key growth factor for lymphatic vessel formation^53^, and VEGF-C is actively involved in brain immunosurveillance^54^. An elevated expression of VEGF-C has been associated with increased lymphatic formation and/or flow^54,100^. Therefore, it is possible that, through the release of vascular mediators, mast cells directly modulate meningeal lymphatic vasculature function^101^. The sagittal sinus is also an area where CNS-derived antigens accumulate and are presented to patrolling T-cells^87^. There is substantial *in vivo* and *in vitro* evidence demonstrating that mast cells can modulate T-cells’ activity^102,103^, and a recent study showed specifically that meningeal mast cells play a crucial role in T-cell recruitment and induction of pathogenesis in a model of experimental encephalomyelitis^104^. Taken together, an increased expression of pseudopodia in the sagittal sinus could indicate an enhanced interaction between mast cells and lymphatic vessels and/or T-cells, facilitating immune trafficking.

### Early life adversity exerts sex-specific effects on meningeal inflammatory markers

To assess whether ELA has long-term effects on meningeal immunology, we performed gene expression analysis in whole duramater samples of adult males and females exposed to NMSEW. We found that, while Tph1 and Hdc were not affected by early life treatment, CMA1, IL-6, and TNFαwere elevated in females but not males exposed to NMSEW. These results suggest that females may exhibt greater ELA-induced meningeal inflammation, which is in line with a wealth of research demonstrating that women show stronger inflammatory responses^105–107^ and heightened susceptibility to diseases associated with ELA including anxiety and depression^108^, migraines^109,110^, and autoimmune disorders^111^. The present study provides evidence that this exaggerated NMSEW-induced dural inflammation in females is mediated, at least in part, by mast cell activity. First, we found that inhibition of mast cell function by administration of ketotifen was sufficient to prevent the proinflammatory effects of ELA. Second, our histological analyses showed that, compared to males, females have elevated activity of mast cells located in the dural transverse sinus, a key neuroimmune interface^87^ as discussed above. Previous work showed that mast cells from female mice have an increased baseline mediator storage capacity and show exaggerated mediator release in response to psychological stress compared to mast cells from males^112,113^. Since ELA exerts long term effects on mast cell function^50,114^, it is possible that a female-biased mast cell reactivity results in stronger proinflammatory effects. Whether there is a causal relationship between increased meningeal inflammation and persistent, low-grade systemic inflammation contributes to ELA-related disease risk remains to be elucidated.

In sum, our studies show that ELA can have long-lasting, sex-specific effects on meningeal function and suggest that mast cells could play a critical role connecting the central nervous system with peripheral immune responses. Since meninges are a crucial interface between the brain and peripheral immunity, these results raise the intriguing hypothesis that meningeal mast cells could be directly contributing to the enhanced crosstalk between stress-processingbrain networks and the immune system thought to underly ELA-induced vulnerability to CNS disease^27,28,87^. A better understanding of the role of meningeal mast cells in CNS and systemic inflammation could provide key molecular insights aiding development of novel, mast cell-targeting therapeutics for stress-related diseases.

## Acknowledgements

The authors would like to thank Kyan Thelen and Ken Moon for their excellent technical assistance. This work was supported by a Brain and Behavior Research Foundation 2019 NARSAD Young Investigator Award to NDW; R01 HD072968 to AJM and AJR.

